# Synergistic Therapy of Doxorubicin with Cationic Anticancer Peptide L-K6 Reverses Multidrug Resistance in Cancer Cells in vitro via P-glycoprotein Inhibition

**DOI:** 10.1101/2021.02.02.429308

**Authors:** Che Wang, Lili Huang, Ruojin Li, Ying Wang, Xiaoxue Wu, Dejing Shang

## Abstract

Multidrug resistance (MDR) is one of the major obstacles to efficient chemotherapy against cancers, resulting from the overexpression of drug efflux transporters such as P-glycoprotein (P-gP). In the present study, we aimed to evaluate the MDR reversal activity and synergistic therapeutic potential of cationic anticancer peptide L-K6 with doxorubicin (DOX) on P-gP-overexpressing and DOX-resistant MCF-7/Adr human breast cancer cells. Flow cytometry and confocal laser scanning microscopy were used to determine the intracellular accumulation of DOX and another P-gP substrate, Rho123. P-gP-Glo assay, Western blot and Biacore analysis were further performed to evaluate the P-gP function and expression. The cytotoxicity in MCF-7 or MCF-7/Adr cells was measured by MTT assay. Flow cytometry assay and confocal laser scanning microscopy observation clearly revealed an increased intracellular accumulation of DOX and Rho123 in MCF-7/Adr cells treated with L-K6, suggesting a P-gP inhibiting potential. Biacore analysis, P-gP-Glo assay and Western blot further confirmed that L-K6 could directly interact with P-gP, inhibit P-gP function and decrease P-gP expression in MCF-7/Adr cells. In addition, as expected, the data from MTT assay indicated that L-K6 restored the sensitivity of MCF-7/Adr cells to DOX, indicating a MDR reversal potential and a promising synergistic anticancer activity. All these findings may provide experimental evidence to support the promising applications and synergistic therapeutic potential of peptidic P-gP inhibitors against MDR cancer.

## 1. Introduction

Multidrug resistance (MDR) is the mechanism by which cancers develop resistance to numerous chemotherapeutics, thus significantly decreases the therapeutic efficacy of clinical anticancer drugs. The most common cause for cancer MDR is the overexpression of various cell membrane-bound ATP-binding cassette (ABC) transporters. These proteins were coupled with energy-consumption pump to efflux endogenous cellular metabolites or exogenous toxic substances out of cells. However, in MDR cancer cells, these proteins can exocytose various chemotherapeutics and in turn attenuate their anticancer effects. As important members of ABC transporter family, P-glycoprotein (P-gP), breast cancer resistance protein (BCRP), and multidrug resistance-associated protein 1 (MRP1) are all responsible for cancer MDR [1]. The inhibition of these ABC transporter proteins may help reverse MDR and restore the attenuated anticancer activities of conventional chemotherapeutics [2–4].

Among these MDR-related transporter proteins, P-gP is the most prominent one and has been well-characterized in many MDR cancer cells [5,6]. P-gp is a 170-180 kDa membrane glycoprotein encoded by MDR1 gene and is first discovered in drug-resistant Chinese hamster ovarian cells [7]. As an ATP-dependent drug efflux pump, P-gP exocytoses various anticancer drugs, such as vincristine, doxorubicin (DOX), and actinomycin D, out of cancer cells [8,9]. Despite anticancer drugs, P-gP also transports phosphatidylserine (PS), a negative charged lipid composition of cell membrane, from the inner to the outer leaflet of the plasma membrane, and thus increases the net negative charge on cell surface, which may provide potential target for cancer specific therapy. Previous studies including ours have found that various cancer cells have an elevated level of cell surface PS [10–12] and exert a higher susceptibility to cationic peptides than noncancerous normal cells [12–14].

In this study, we screened the P-gP inhibiting activity of various cationic peptides and found L-K6, one lysine-rich peptide, significantly inhibited P-gP function and reduced P-gP expression in DOX-resistant MCF-7/Adr cells. Moreover, we further conducted a series of experiments to investigate the reversal effect of L-K6 on P-gP-mediated MDR. We also analyzed the synergistic potential of L-K6 with DOX in MCF-7/Adr cells and hope to explore the possible roles of L-K6 in the combination therapy against MDR breast cancers.

## 2. Materials and Methods

### 2.1 Reagents and antibody

L-K6 was synthesized and purified (>95%) according to our previously described protocol [15,16]. L-K6 was dissolved in saline and further diluted in culture medium to required concentrations. Verapamil (VER) and DOX (Sigma-Aldrich Corp., St. Louis, MO) were dissolved in dimethyl sulphoxide (DMSO) and further diluted in culture medium with the final DMSO concentration < 0.1%. Monoclonal antibody against P-gP was obtained from Abcam (Cambridge, UK). RPMI Medium 1640 and fetal bovine serum (FBS) were obtained from GIBCO (Grand Island, NY, USA). The 3-(4,5-dimethylthiazol-2-yl)-2,5-diphenyl-2H-tetrazolium bromide (MTT, purity > 98%) was purchased from Sigma-Aldrich (St. Louis, MO, USA).

### 2.2 Cell culture

DOX-sensitive human breast cancer cell line MCF-7 and its DOX-resistant derivative P-gP-overexpressing MCF-7/Adr cells were purchased from Nanjing KeyGen Biotech Co. Ltd. (Nanjing, China). Cells were cultured in RPMI 1640 supplemented with 10% FBS, 1% non-essential amino acid solution, 100 U/mL penicillin and 0.1 mg/mL streptomycin. All cells were cultured at 37 °C in a humidified atmosphere of 5% CO_2_.

### 2.3 MTT assay

MTT assay was performed to assess the viability and sensitivity of cells to DOX with or without various concentrations of L-K6. Both MCF-7 and MCF-7/Adr cells were harvested, resuspended and seeded in 96-well plates (100 μL aliquots, final density at 4×10^5^ cells/mL). After 24 h of culture, cells were exposed to a series of concentrations of DOX or L-K6 alone, or a combination of L-K6 (10-50 μM) with DOX (10-120 μM), and were cultured at 37 °C for 24 h. Then, MTT (5 mg/mL, 20 μL/well) was added to each well and cells were incubated for further 4 h at 37 °C, allowing the viable cells to reduce the yellow MTT into purple-blue formazan crystals. After adding 100 μL/well of DMSO, optical density values were measured at 570 nm with a microplate reader (Bio-Rad, San Diego, CA, USA). Six wells were used for each drug concentration and the experiment was repeated 3 times. The IC_50_ value was calculated by Prism (Graphpad Software, La Jolla, CA, USA) [17]. The resistance index (RI) was calculated by dividing the IC_50_ values of the MCF-7/Adr cells by that of the MCF-7 cells. The fold-reversal factor (Fr) of MDR was calculated by dividing the IC_50_ of DOX in the absence of L-K6 by those obtained in the presence of L-K6. The synergistic index (Q) was calculated by using following formula, in which E presents the percentage of proliferation inhibition. Q=E_(DOX+L-K6)_/(E_DOX_+E_L-K6_-E_DOX_×E_L-K6_)

### 2.4 Doxorubicin accumulation assay by flow cytometry

DOX accumulation was determined using the protocol as previously described with minor modifications [18]. Briefly, MDR cancer cells were seeded in 6-well plates. After 24 h incubation of the cells with L-K6, DOX was added and incubated for 1 h at 37 °C. Cells were then washed with phosphate buffered saline (PBS), trypsinized and resuspended in medium (10^6^ cells/mL) [19]. The intracellular auto-fluorescence of DOX was determined by flow cytometry (FACSCalibar, BD, USA) with excitation measured at 488 nm and emission measured at 575 nm.

### 2.5 Doxorubicin accumulation assay by confocal laser scanning microscopy

Coverslips were autoclaved and placed into the wells of a 6-well plate. MCF-7/Adr cells were seeded onto the coverslips. After 48 h, cells were washed with PBS for three times and treated with DOX for 1 hour, then were fixed with 2% (w/v) paraformaldehyde in PBS. The coverslips were wet-mounted on microscope slides and observed under the Zeiss LSM 710 confocal laser scanning microscope (Jena, Germany) with Plan-Neofluar 40X/1.3 Oil DIC lens to determine the intracellular DOX accumulation as described previously [20].

### 2.6 Rho123 accumulation determined by flow cytometry

The intracellular accumulation of specific P-gp substrate Rho123 was determined by flow cytometry according to standard procedures [21]. The cells treated with L-K6 were cultured at 37°C for 24 h, and then incubated for 0.5□h with 5□μg/mL Rho123. The cells were harvested and washed three times with cold PBS. The fluorescence intensity was analyzed by flow cytometry (FACSCalibar, BD, USA).

### 2.7 Biacore Surface Plasmon Resonance Spectroscopy analysis

Surface plasmon resonance spectroscopy analysis was performed by using research grade CM5 sensor chip (Biacore, Inc. USA). The P-gP monoclonal antibody was firstly immobilized on CM5 chip by using amine coupling kit (Biacore, Inc. USA). The antibody-immobilized chip was then conditioned by passing membrane protein extraction of MCF-7/Adr cells to capture P-gP proteins. The L-K6 peptides were diluted in HEPES Buffered Saline (10 mM HEPES with 3 mM EDTA, 0.005% surfactant P20, and 150 mM NaCl, pH 7.4). BSA (0.1 mg/ml) was used as negative control. The prepared 30 μL of L-K6 flowed over the P-gP-immobilized chip at a rate of 20 μL/min, and the association and dissociation were measured.

### 2.8 P-gP ATPase activity assay

Stimulation of P-gP ATPase activity was utilized to indicate the interaction between peptides and P-gP using the P-gP-Glo Assay System (Promega, USA) according to the manufacturer’s instructions. Sodium orthovanadate (Na_3_VO_4_) was used as a P-gP ATPase inhibitor. The P-gP inhibitory effects of L-K6 were determined against a VER-stimulated P-gP ATPase activity. Various concentrations of L-K6 were incubated in 100 μM VER, 5 mM MgATP and 25 μg recombinant human P-gP membranes at 37 °C for 40 min. Luminescence was initiated by ATP detection buffer. After incubation at room temperature for 20 min to allow luminescence development, the untreated white opaque 96-well microplates were read on luminometer (SpectraMax M5, Molecular Devices, USA). The changes of relative light units (ΔRLU) were determined by comparing Na_3_VO_4_-treated samples with L-K6/VER-treated group.

### 2.9 Western blotting assay

As previously described [22], cells were homogenized on ice in lysis buffer for Western blot. Proteins from each cell lysate were mixed with loading buffer (125 mM Tris-HCl, pH 6.8, 4% sodium dodecyl sulfate, 20% glycerol, 10% β-mercaptoethanol, and 0.05% bromophenol blue). Equal amounts of proteins were loaded and separated in SDS-PAGE by electrophoresis. Proteins were transferred onto PVDF membranes by electroblotting. Membranes were blocked with 5% BSA in Tris-buffered saline with 0.1% Tween 20 for 2 h at 37 °C, and then incubated with antibody overnight at 4 °C. Finally, blots were developed with horseradish peroxidase-conjugated anti-IgG antibody and detected by an enhanced chemiluminescence (ECL) method using ECL Western Blotting Detection system (Amersham Biosciences, UK). The intensities of the protein bands were quantified with densitometry using Scion Image software.

### 2.10 Statistical analysis

Data are expressed as means ± SD. All experiments were repeated at least 3 times and the differences were determined by using Student’s t test. The value of *p* < 0.05 was considered to be statistically significant.

## 3. Results

### 3.1 L-K6 significantly altered the expression level of P-gP in MCF-7/Adr cells

To validate the overexpression of P-gP in MCF-7/Adr cells, the P-gP levels of MCF-7 and MCF-7/Adr cells were determined and compared by Western blotting analysis. As shown in supplementary Fig. S1, MCF-7/Adr cells showed a significantly higher level of P-gP expression than its parental MCF-7 cells.

To further evaluate the inhibiting effects of peptides on P-gP expression, Western blotting analysis was performed in L-K6-treated MCF-7/Adr cells. As shown in Fig. 1, L-K6 significantly reduced P-gP expression in MCF-7/Adr cells in a dose-dependent manner (Fig. 1A and 1B).

**Figure 1.**
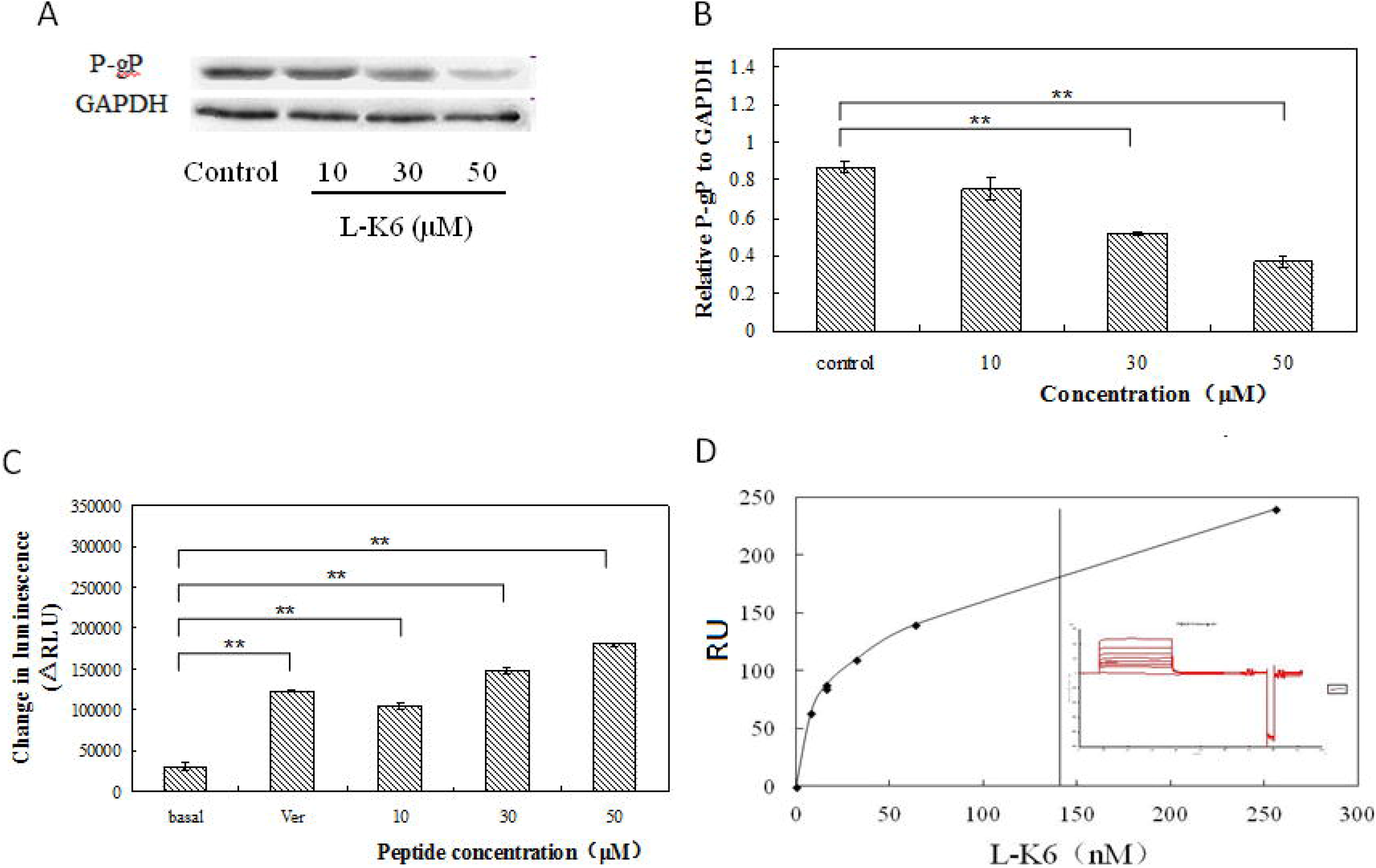
Dual Effects of L-K6 on P-gp pump and expression. (A) after 24 h exposure, L-K6 (10-50 μM) dose-dependently reduced P-gp expression. (B) The protein expression of P-gp was normalized to β-actin. (C) L-K6 significantly inhibited P-gP-ATP activity as assessed by P-gP-GLO assay; (D) Direct interactions of L-K6 with P-gP protein as assessed by SPR. Each column shows the mean ± SD of 3 independent experiments, performed in triplicate. *P < 0.05 and **P < 0.01, vs. the control group.

### 3.2 L-K6 significantly inhibited P-gP-ATP activity

Despite the reduction of P-gP expression, we further assess the inhibiting activity of peptides on P-gP function by P-gP-GLO assay, using non-peptidic P-gP inhibitor VER as a positive control. As shown in Fig. 1C, VER strongly induced ATP accumulation (corresponding to its P-gP ATPase inhibiting activity), which was dose-dependently enhanced also by L-K6, indicating a direct P-gP inhibiting activity of L-K6.

### 3.3 Direct interactions of L-K6 with P-gP protein as assessed by BIAcore surface plasmon resonance detector

To further confirm the direct interactions of L-K6 with P-gP, the BIAcore surface plasmon resonance (SPR) assay was performed in vitro. As shown in Fig. 1D, the data from SPR clearly demonstrated that L-K6 peptide could tightly interact with P-gP protein with a KD value of 141 nmol.

### 3.4 L-K6 increased the intracellular DOX accumulation in MCF-7/Adr cells

To further confirm the inhibiting effect of L-K6 on P-gP function, we determined the intracellular accumulation of DOX in L-K6-treated MCF-7/Adr cells. The FACs analysis clearly revealed that L-K6 could significantly increase the intracellular DOX accumulation in MCF-7/Adr cells, which was comparable to the positive control VER (Fig. 2). L-K6 at 10, 30 and 50 μM concentrations increased the intracellular accumulation of DOX by 1.27-, 1.48- (*p* <0.01) and 1.19-fold (*p* <0.01) in MCF-7/Adr cells, respectively. This DOX-efflux inhibiting effect could be also observed by confocal laser scanning microscopy, as shown by the increased red-fluorescence density (Fig. 3). All these results suggested that L-K6 significantly inhibited DOX efflux in MCF-7/Adr cells.

**Figure 2.**
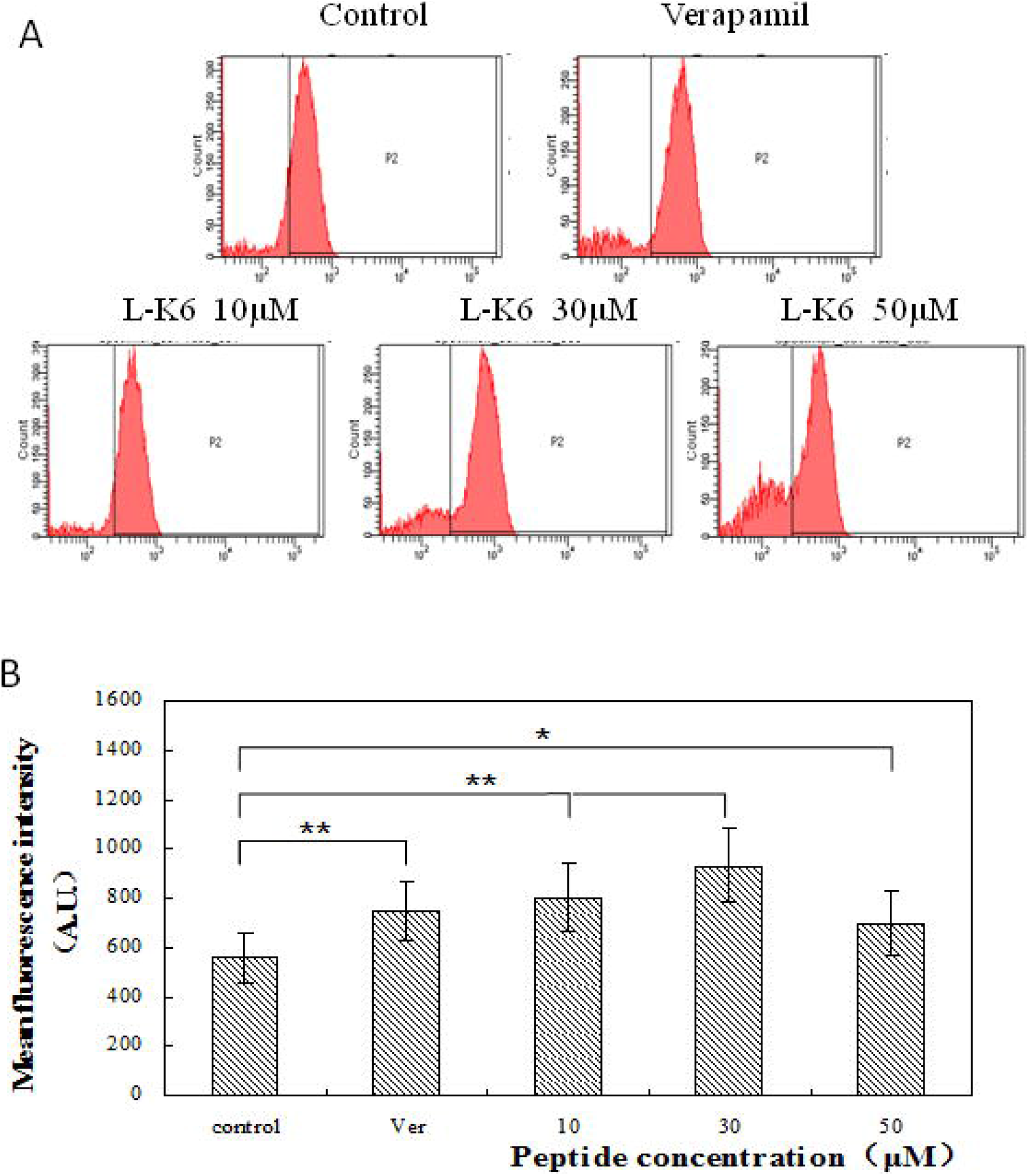
Effects of L-K6 on intracellular accumulation of DOX in MCF-7/Adr cells. (A) Cells were pre-incubated with or without L-K6 or verapamil for 24 h at 37°C and then incubated with 20 μM DOX for another 1h at 37°C. The intracellular auto-fluorescence of DOX was determined by flow cytometric analysis. (B) The results are presented as fold change in fluorescence intensity relative to untreated control resistant cells. Data shown are means ± SD of triplicate determinations. *P < 0.05 and **P < 0.01 vs. the control group.

**Figure 3.**
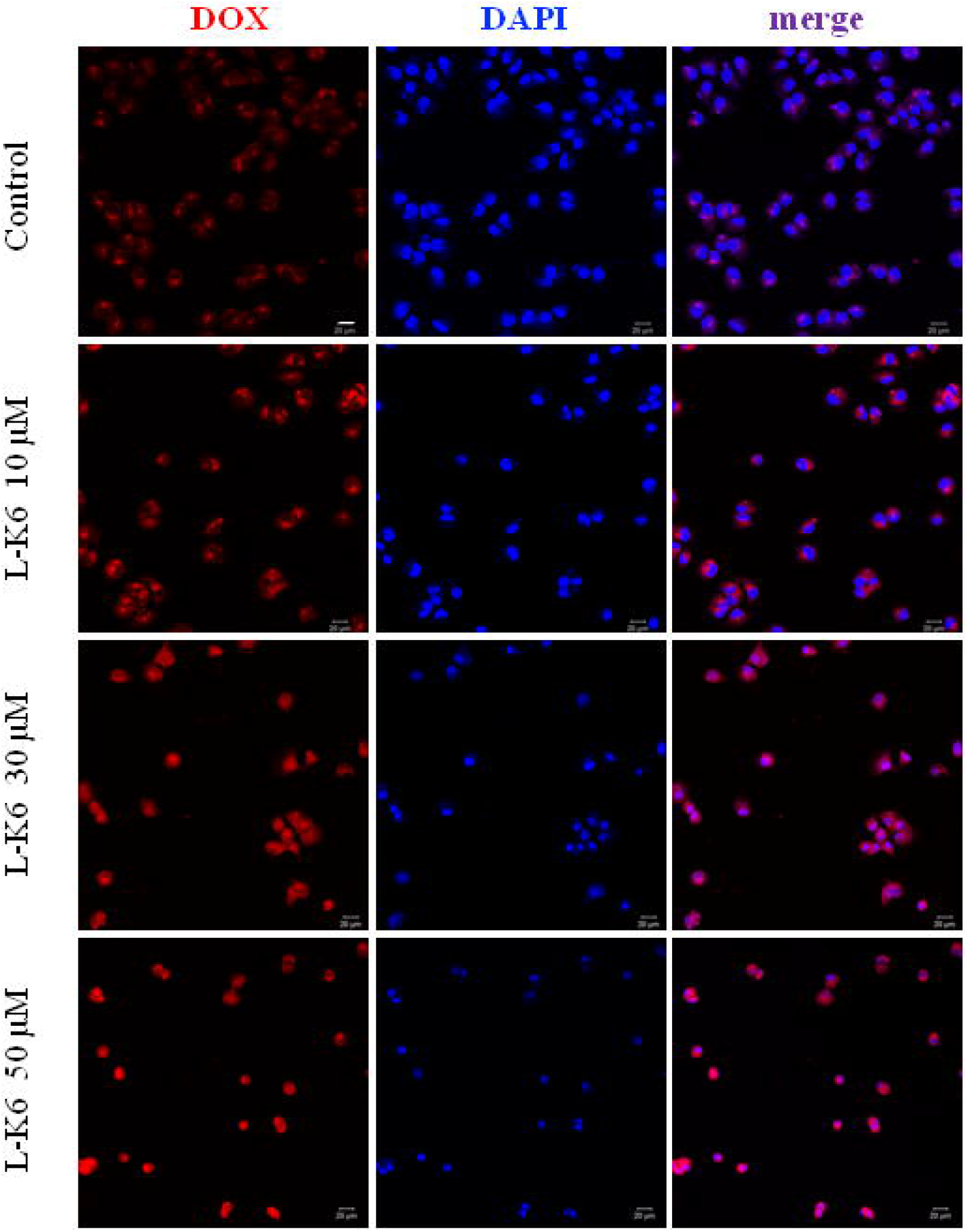
Effects of L-K6 on intracellular accumulation of DOX in MCF-7/Adr cells. Cells were pre-incubated with or without L-K6 for 24 h at 37°C and then incubated with 20 μM DOX for another 1h at 37°C. The intracellular auto-fluorescence of DOX was observed by confocal microscopy.

Interestingly, our data also indicated that at higher concentration (50 μM), the level of intracellular DOX accumulation was decreased. This might be due to the slight increase of membrane permeability (supplementary Fig. S2).

### 3.5 L-K6 increased the intracellular Rho123 accumulation in MCF-7/Adr cells

To further confirm the P-gP substrate potential and the reversal activity of L-K6 on P-gP-mediated drugs efflux, the intracellular accumulation of another P-gP substrate, Rho123, was also determined by flow cytometry. Our data clearly demonstrated that L-K6 also significantly increased the intracellular Rho123 fluorescence density in MCF-7/Adr cells, which was comparable with P-gP inhibitor VER (Fig. 4).

**Figure 4.**
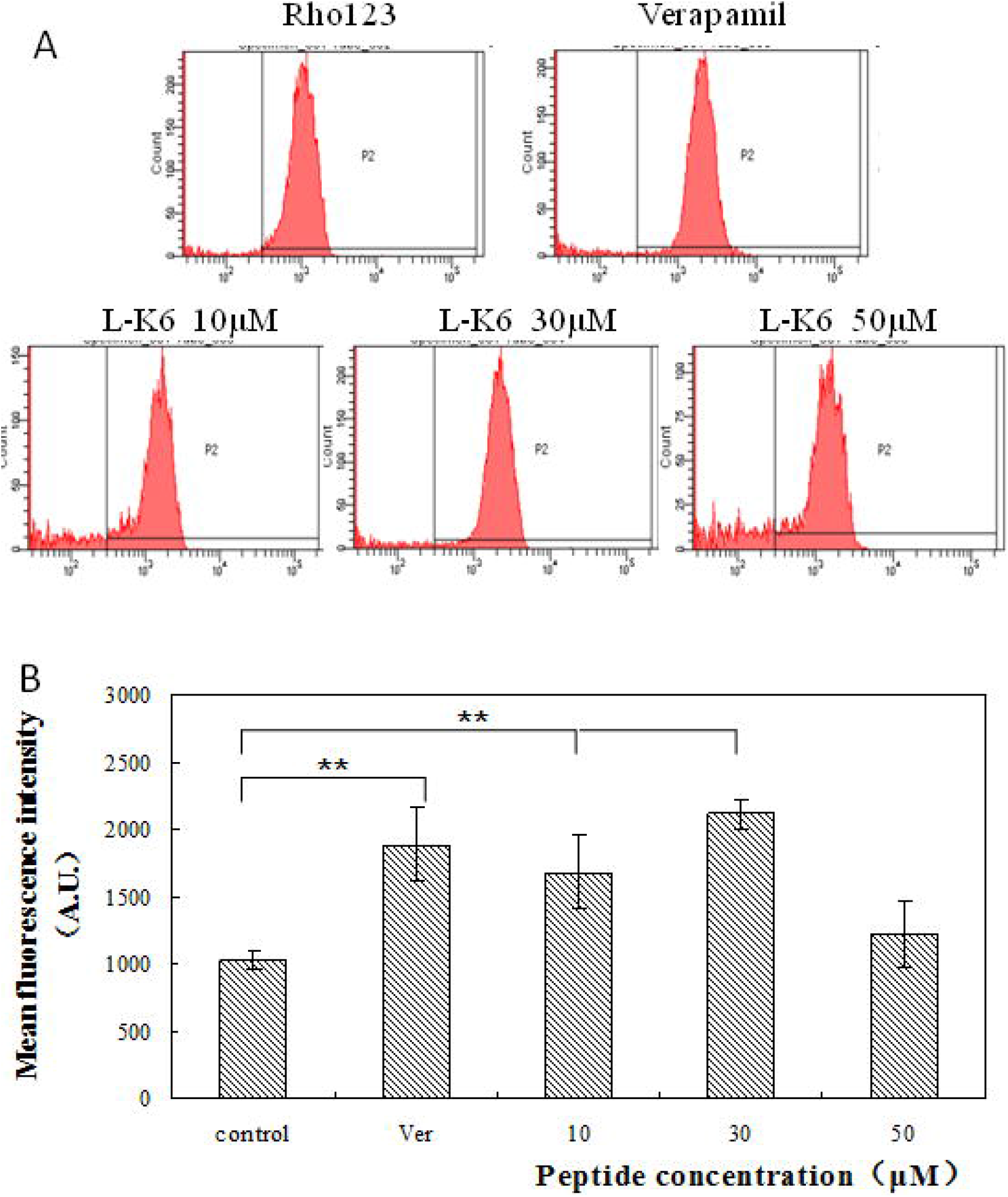
Effects of L-K6 on intracellular accumulation of Rho123 in MCF-7/Adr cells. (A) Cells were pre-incubated with or without L-K6 or verapamil for 24 h at 37°C and then incubated with Rho123 for another 1h at 37°C. The intracellular fluorescence of Rho123 was determined by flow cytometric analysis. (B) The results are presented as fold change in fluorescence intensity relative to untreated control resistant cells. Data shown are means ± SD of triplicate determinations. *P < 0.05 and **P < 0.01 vs. the control group.

Consistent with the data from DOX accumulation assay, we found that the level of intracellular Rho123 accumulation was also decreased after higher concentration (50 μM) of L-K6 exposure (Fig. 4).

### 3.6 Cytotoxic responses of MCF-7 and MCF-7/Adr cells to DOX

The MTT data indicated a resistant response of MCF-7/Adr cells to DOX-induced cytotoxicity, compared with parental MCF-7 cells (Fig.5A). After 24-h DOX exposure, a dramatic cell death was observed in MCF-7 cells. The IC_50_ value of DOX is 5.3 ± 0.4 μM, indicating a good sensitivity of MCF-7 cancer cells to DOX. In contrast, the P-gP-overexpressed MCF-7/Adr cells showed a significant resistance to cytotoxicity induced by DOX, evidenced by the dramatic increase of IC_50_ value (258.3 ± 10.9 μM vs. 5.3 ± 0.4 μM) (Fig. 5A). The resistance index (RI) value of MCF-7/Adr cells to DOX is 48.7, compare with its parental cell line MCF-7 (Fig. 5A).

**Figure 5.**
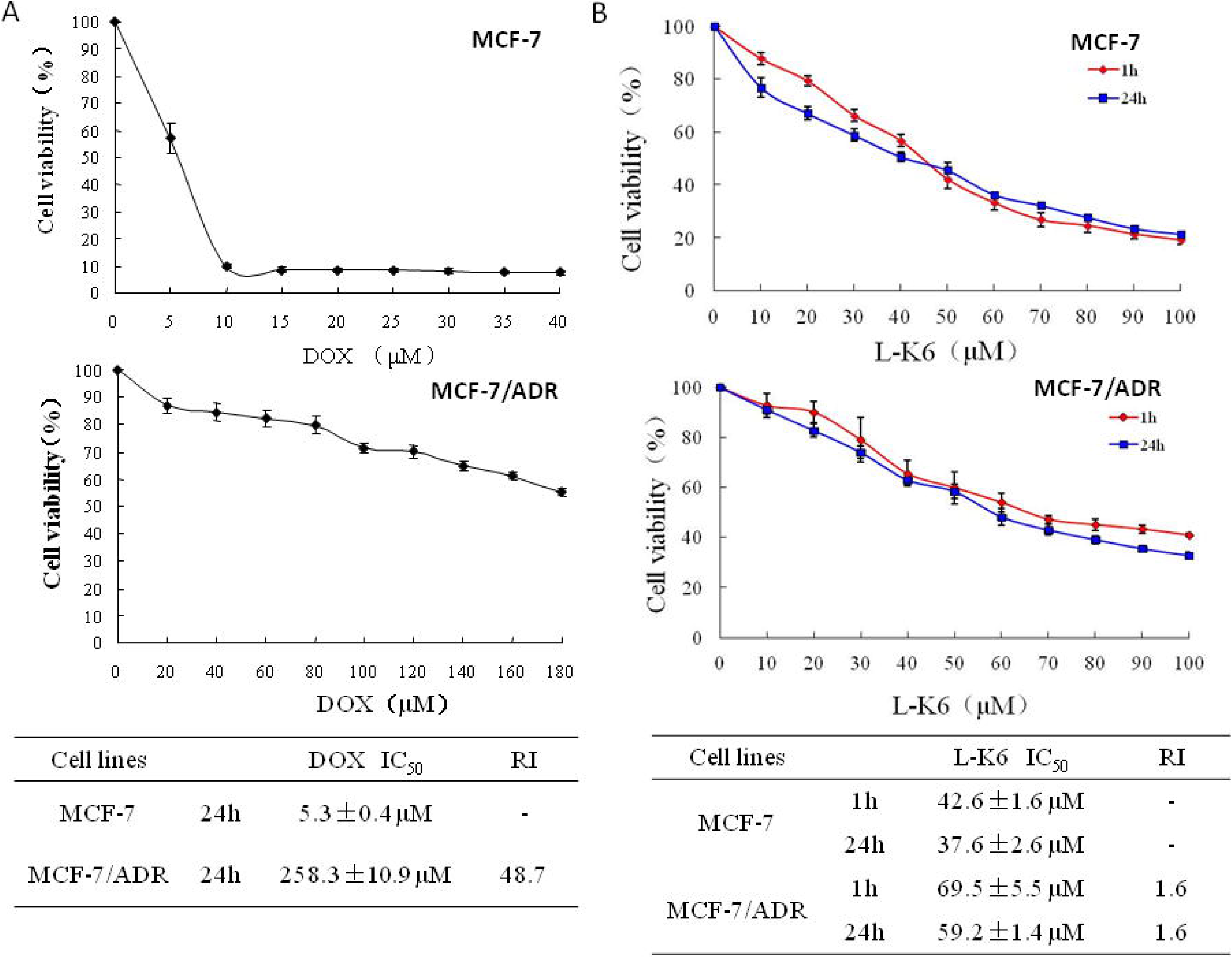
Cytotoxic effects of DOX and L-K6 in MCF-7 and MCF-7/Adr cells. (A) The MTT assay and IC50 values of DOX in MCF-7 and MCF-7/Adr cells; (B) The MTT assay and IC50 values of L-K6 in MCF-7 and MCF-7/Adr cells; RI, the resistance index of each drug.

### 3.7 Cytotoxic effects of L-K6 in MCF-7 and MCF-7/Adr cells

Consistent with our previous findings [15], in the present study, cationic L-K6 peptide induced a rapid cytotoxicity in MCF-7 cells in a dose-dependent manner, with IC_50_ values of 42.6 ± 1.6 (1 h) and 37.6 ± 2.6 (24 h) (Fig. 5B). Interestingly, the data from our MTT assay in the present study unexpectedly indicated that MCF-7/Adr cells also showed a resistance to anticancer peptide L-K6, evidenced by the slightly increased IC_50_ values compared with MCF-7 cells (69.5 ± 5.5 vs. 42.6 ± 1.6 μM at 1 h, and 59.2 ± 1.4 vs. 37.6 ± 2.6 μM at 24 h, respectively, *p* <0.05). The RI values were both 1.6 at 1 and 24 h time points (Fig. 5B).

### 3.8 L-K6 partially reversed the MDR in MCF-7/Adr cells to DOX

To test the possible MDR reversal effect of L-K6 to MCF-7/Adr cells against DOX, various concentrations of L-K6 (10-50 μM) was used in reversal assay. As shown in Fig. 6A, the co-administration of L-K6 partially reversed the MDR response of MCF-7/Adr cells to DOX in a dose-dependent manner. IC_50_ values of DOX in MCF-7/Adr cells were significantly decreased after L-K6 exposure (Fig.6B). The fold-reversal factor (Fr) values of MDR were 2.65, 3.24 and 3.84 in the presence of L-K6 at 10, 30 and 50 μM concentrations in MCF-7/Adr cells, respectively (Fig. 6B).

**Figure 6.**
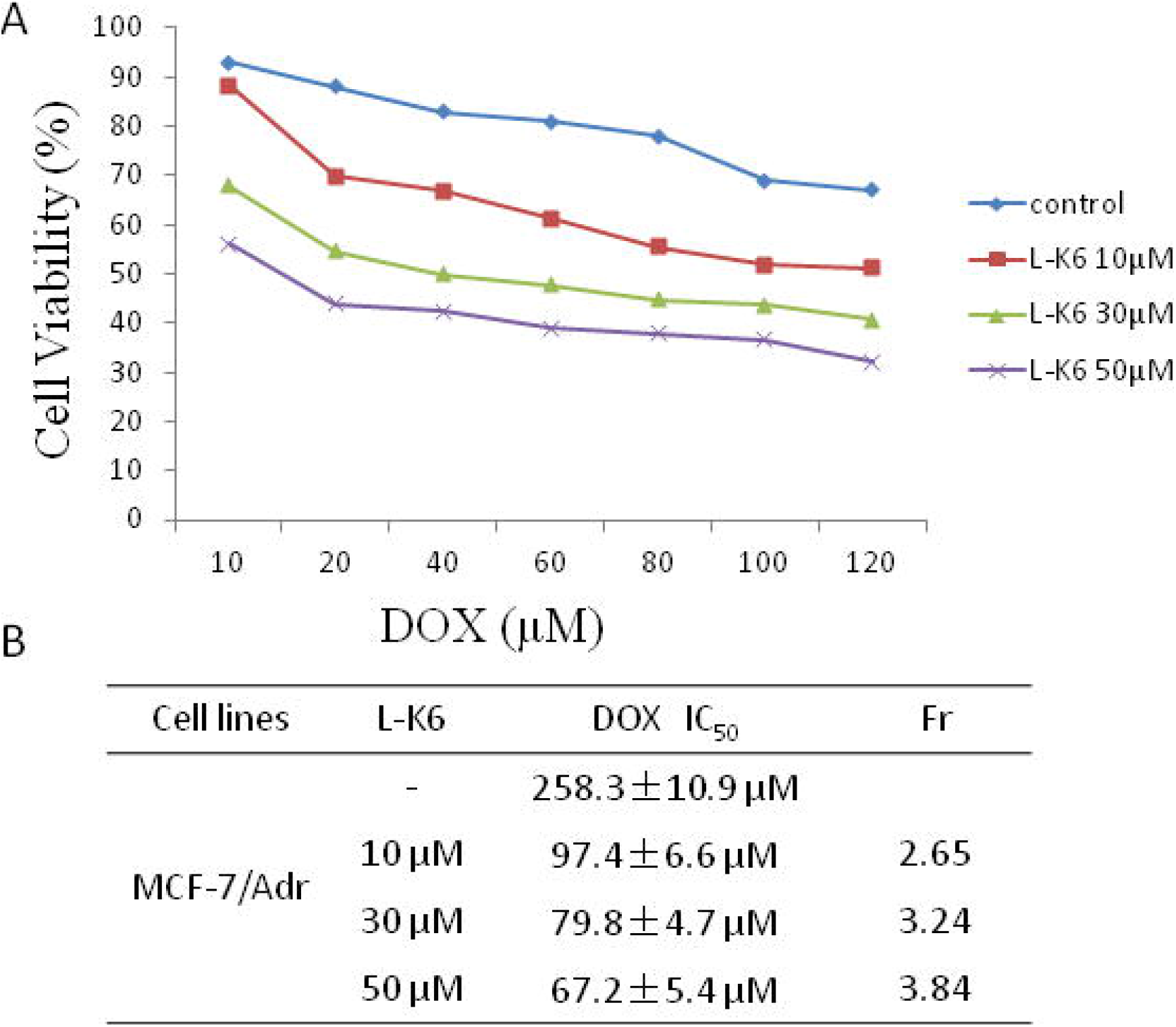
Combined treatment of L-K6 and DOX reverses MDR in MCF-7/Adr cells. (A) combined treatment with L-K6 and DOX dose-dependently decreased the cells viability of MCF-7/Adr cells; (B) L-k6 showed additive and synergistic activity to DOX.

L-K6 at as low as 10 μM (non-lethal) concentration could significantly decrease the IC_50_ value of DOX in MCF-7/Adr cells (258.3 ± 10.9 vs. 97.4 ± 6.6 μM, Fig. 6B), while at this concentration L-K6 caused only minimal cytotoxicity to MCF-7/Adr cells (cell viability was over 90%, Fig. 5B). Although the IC_50_ values of DOX plus L-K6 treatment in MCF-7/Adr cells is still higher than those of DOX-treated MCF-7 cells (97.4 ± 6.6, 79.8 ± 4.7, 67.2 ± 5.4 vs. 5.3 ± 0.4 μM), our data suggested that L-K6 could partially reverse the MDR of P-gp-overexpressing MCF-7/Adr cells to DOX exposure.

### 3.9 Synergistic activity of L-K6 on DOX-induced cytotoxicity in MCF-7/Adr cells

Much more importantly, further data analysis revealed an additive or synergistic activity of L-K6 with DOX in MCF-7/Adr cells. Normally, when 0.85□<□Q□<□1.15, the effect of combined therapy is additive; when Q □ 1.15, the effect of combined therapy is synergistic [23]. As summarized in Table 1, our data suggested that L-K6 at 10 and 50 μM showed an additive effect with DOX, whereas L-K6 at 30 μM presented synergistic activity with DOX (40~100 μM) in MCF-7/Adr cells (Table 1).

**Table 1.** Synergistic anticancer activity (Q values) of L-K6 with DOX

| | | DOX ($\mu\text{M}$) | | | | | |
| --- | --- | --- | --- | --- | --- | --- | --- |
|  |  | 20 | 40 | 60 | 80 | 100 | 120 |
| L-K6<br>($\mu\text{M}$) | 10 | 0.92 | 0.95 | 1.23 | 1.14 | 1.04 | 0.9 |
|  | 30 | 1.11 | 1.19 | 1.21 | 1.19 | 1.17 | 1.12 |
|  | 50 | 1.1 | 1.11 | 1.15 | 1.12 | 1.11 | 1.1 |

## 4. Discussion

Breast cancer is the first common cancer and the second leading cause of cancer death among females, next only to lung cancer [24]. Despite the significant advances in the diagnosis and chemical or biological therapies of breast cancer, metastatic breast cancer is still incurable due to the development of rapid resistance [25,26]. Therefore, identification of novel candidates with MDR reversal activity will help improve the clinical benefits of current conventional anticancer therapeutics. P-gP, as an important member of ABC superfamily and a multi-specific drug efflux transporter, plays a significant role in governing the bioavailability of many clinically active drugs. The inhibition of P-gP function or expression by various inhibitors forms a distinctive approach in improving bioavailability and conquering drug resistance.

In the present study, MCF-7/Adr cells were verified to be DOX-resistant (Fig. 5), via mechanisms including the overexpression of P-gP protein (supplementary Fig. S1). Our data also indicated that L-K6 at reversal concentrations (10-50 μM) remarkably potentiated the cytotoxicity of DOX in P-gP-overexpressing MCF-7/Adr cells (Fig. 6). Additionally, the combined therapy of L-K6 (30 μM) with DOX (40-100 μM) showed a synergistic activity, evidenced by Q values over 1.15.

The MDR-reversal and synergistic activity of L-K6 in MCF-7/Adr cells might be due to the potential impacts of L-K6 on P-gP function and expression. Our data indicated that L-K6 could significantly increase the intracellular accumulation of P-gP substrates DOX and Rho123 in MCF-7/Adr cells (Fig. 2 and 4), predicting a P-gP inhibiting potential of L-K6. Biacore SPR assay further revealed a direct interaction between L-K6 and P-gP protein. Moreover, the P-gP-GLO assay further suggested that L-K6 could interact with P-gP and inhibit P-gP ATPase activity (Fig. 5C and 5D), thereafter depressed the P-gP function to efflux bioactive substrates.

Moreover, the overexpressed P-gP in MDR MCF-7/Adr cells should theoretically lead to an elevated negatively charged PS on cell surface and consequently a higher affinity of MDR cells membrane to the cationic peptides. However, in our present study, MCF7/Adr cells were resistant to L-K6, compared to parental MCF-7 cells. Considering the tight interactions between L-K6 and P-gP in SPR assay, this resistance suggested that L-K6 might be a potential substrate of P-gP and could competitively inhibit the P-gP-mediated efflux of DOX and thus reversed the resistance of MCF-7/Adr cells to DOX-induced cytotoxicity.

Our experiments further suggested that the MDR reversal effect of L-K6 is not only due to the inhibition of P-gp function, but also via the down-regulation of P-gp expression. Inhibition of P-gP expression is an ideal way to overcome cancer MDR. Several compounds have been previously reported to have potency to down-regulate P-gP expression, including the calcium channel blockers VER [27,28], and some natural occurring compounds such as curcuminoids [29,30] and cobalamin [31]. In this study, we found a new peptidic member with activities to down-regulate P-gP in the MDR cancer cells with overexpressed P-gP. Our Western blotting results showed that cationic anticancer peptide L-K6 down-regulated the protein expression level of P-gP in a dose-dependent manner (Fig. 5A and 5B) in MCF-7/Adr cells. It could be suggested that L-K6 reverses P-gp-mediated MDR by both down-regulating P-gp expression and direct inhibiting P-gP-mediated drugs efflux. Although the dual functions of L-K6 inhibiting P-gP’s drug-pump and expression are similar to VER, the cationic nature and anticancer activity may provide a cancer-targeting potential of L-K6 to treat P-gP-overexpressing MDR cancer cells.

## 5. Conclusion

In conclusion, this study provides an evidence that cationic anticancer peptide L-K6 could significantly enhance the efficacy of chemotherapeutic drugs DOX in MDR MCF-7/Adr cancer cells. Furthermore, the reversal of MDR by L-K6 was ascribed to both down-regulation of P-gP expression and direct inhibition of P-gP efflux. Our data further confirmed the MDR reversal ability of L-K6 and may support the potential benefits of combining L-K6 with other conventional antineoplastic agents in overcoming drug resistance in cancer chemotherapy.

## Acknowledgement

This work was supported by the National Natural Science Foundation of China (NSFC 81202448 and 31272314), Scientific Research Fund of Liaoning Provincial Education Department (L2012382, L201683653), the Program for Liaoning Innovative Research Team in University (LT2012019) and Natural Science Foundation of Liaoning Department of Science and Technology (20180550306 and 2019-ZD-0467).

## Author contribution

Che Wang and Dejing Shang proposed the study. Lili Huang, Ruojin Li, Ying Wang and Xiaoxue Wu performed experiments. Che Wang and Lili Huang performed statistical analysis. Che Wang drafted the manuscript.

