## Supplementary figures and images for "Synergistic Therapy of Doxorubicin with Cationic Anticancer Peptide L-K6 Reverses Multidrug Resistance in Cancer Cells in vitro via P-glycoprotein Inhibition"

### Suppl Fig. S1

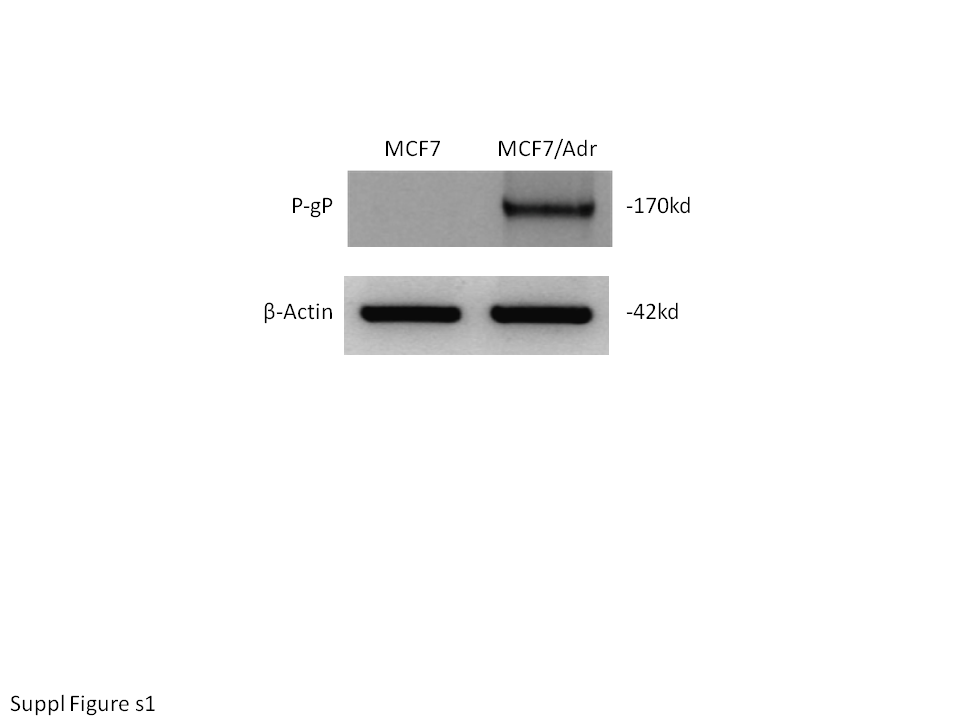

### Suppl Fig. S1

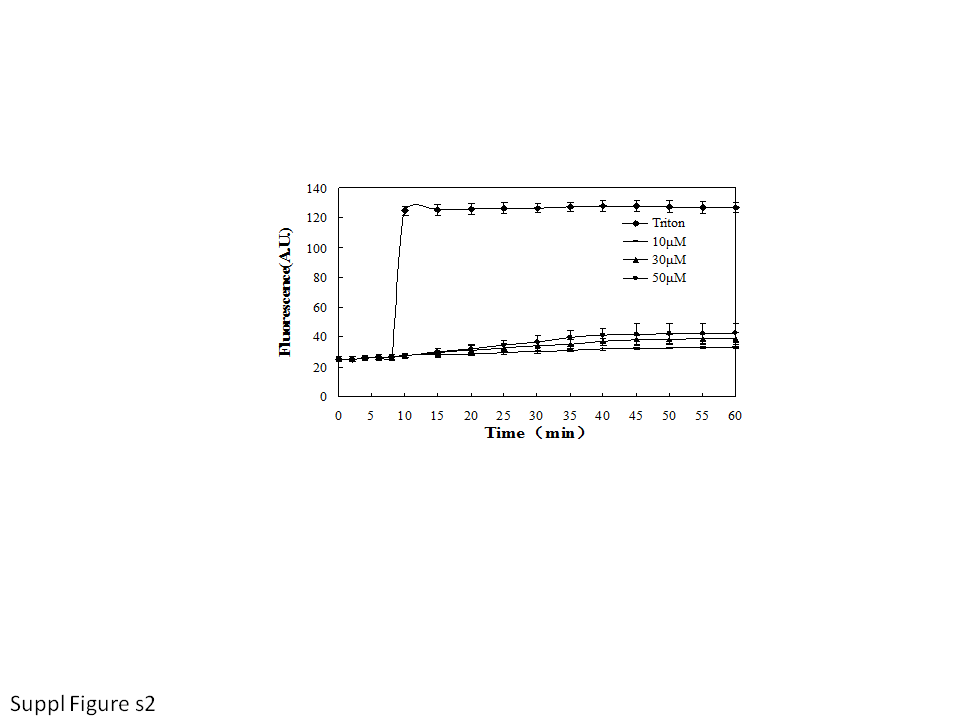
